# The Ontology of Biological Attributes (OBA) - Computational Traits for the Life Sciences

**DOI:** 10.1101/2023.01.26.525742

**Authors:** Ray Stefancsik, James P. Balhoff, Meghan A. Balk, Robyn Ball, Susan M. Bello, Anita R. Caron, Elissa Chessler, Vinicius de Souza, Sarah Gehrke, Melissa Haendel, Laura W. Harris, Nomi L. Harris, Arwa Ibrahim, Sebastian Koehler, Nicolas Matentzoglu, Julie A. McMurry, Christopher J. Mungall, Monica C. Munoz-Torres, Tim Putman, Peter Robinson, Damian Smedley, Elliot Sollis, Anne E Thessen, Nicole Vasilevsky, David O. Walton, David Osumi-Sutherland

## Abstract

Existing phenotype ontologies were originally developed to represent phenotypes that manifest as a character state in relation to a wild-type or other reference. However, these do not include the phenotypic trait or attribute categories required for the annotation of genome-wide association studies (GWAS), Quantitative Trait Loci (QTL) mappings or any population-focused measurable trait data. Moreover, variations in gene expression in response to environmental disturbances even without any genetic alterations can also be associated with particular biological attributes. The integration of trait and biological attribute information with an ever increasing body of chemical, environmental and biological data greatly facilitates computational analyses and it is also highly relevant to biomedical and clinical applications.

The Ontology of Biological Attributes (OBA) is a formalised, species-independent collection of interoperable phenotypic trait categories that is intended to fulfil a data integration role. OBA is a standardised representational framework for observable attributes that are characteristics of biological entities, organisms, or parts of organisms. OBA has a modular design which provides several benefits for users and data integrators, including an automated and meaningful classification of trait terms computed on the basis of logical inferences drawn from domain-specific ontologies for cells, anatomical and other relevant entities. The logical axioms in OBA also provide a previously missing bridge that can computationally link Mendelian phenotypes with GWAS and quantitative traits. The term components in OBA provide semantic links and enable knowledge and data integration across specialised research community boundaries, thereby breaking silos.

## Introduction

Animal models have greatly contributed to the progress of genomics research. In addition to mutant strains identified by traditional phenotypic selection and breeding methods, genome engineering in model organisms allow the generation of transgenic lines and targeted mutants using homologous recombination or CRISPR-Cas9 technology^1–3^. Collectively, these technologies allow researchers worldwide to generate a large body of genetic data using mouse and other model organisms, and the resulting data is made available in several biomedical databases curated by experts^4–7^. Currently there are more than 800 biological databases collecting genotype, phenotype, and variation data from a wide range of organisms^8^. The valuable knowledge in these databases on variants, phenotypes and gene function is highly relevant to human and veterinary medicine, agriculture, evolutionary biology, ecology, and comparative genomics in general.

Technological advances in next-generation DNA sequencing also yield an ever increasing number of new genomic variants with unknown functional significance across the tree of life^9,10^. The identification of the phenotypically and clinically relevant subset of the new DNA variants in both human and veterinary medicine, as well the characterisation of the mechanisms of how these variations exert their phenotypic effects, pose serious challenges that cannot be met successfully without advancements in data integration and computational tools^11^. A standardised and computationally amenable representation of traits is critical for many biomedical and agricultural use cases involving DNA variants, from genome-wide association studies (GWAS) to Quantitative Trait Loci (QTL) mappings^12–16^. Currently, the lack of consistent computational modelling and annotation of traits from various data sources restricts their interoperability and hinders not only genetic mechanisms of discovery for medicine, but also agriculture, biodiversity, and evolutionary biology.

Ontologies provide standardised sets of concepts (terms) that are understandable by human users and also allow for logical inference, computational reasoning, and sophisticated data queries. There are several phenotype ontologies that differ in their scope of specialisation or focus on certain taxonomic groups. For example, the Mammalian Phenotype Ontology (MP)^17^ and the Human Phenotype Ontology (HPO)^18^ have different taxonomic focuses to categorise phenotypes of primarily Mendelian-type inheritance. Each of these ontologies is used to annotate genotypes, where the annotations represent phenotypic states that deviate from a reference, which is usually the wild-type or typical phenotype for the species and population of focus. The phenotypic deviation or abnormality is always represented in the logical axioms in these phenotype ontologies. This is in contrast to trait ontologies, where the logical axioms define generic attributes without reference to any specific phenotypic alterations or states. For example, a “blood lysine amount” can manifest in a “Hypolysinemia” phenotype, where the former manifests in a “decreased amount” phenotypic state with an “abnormal” quality component in the logical equivalence axioms (see Figure 1). This is a fundamental difference between modelling traits and phenotypes ontologically.

**Figure 1:**
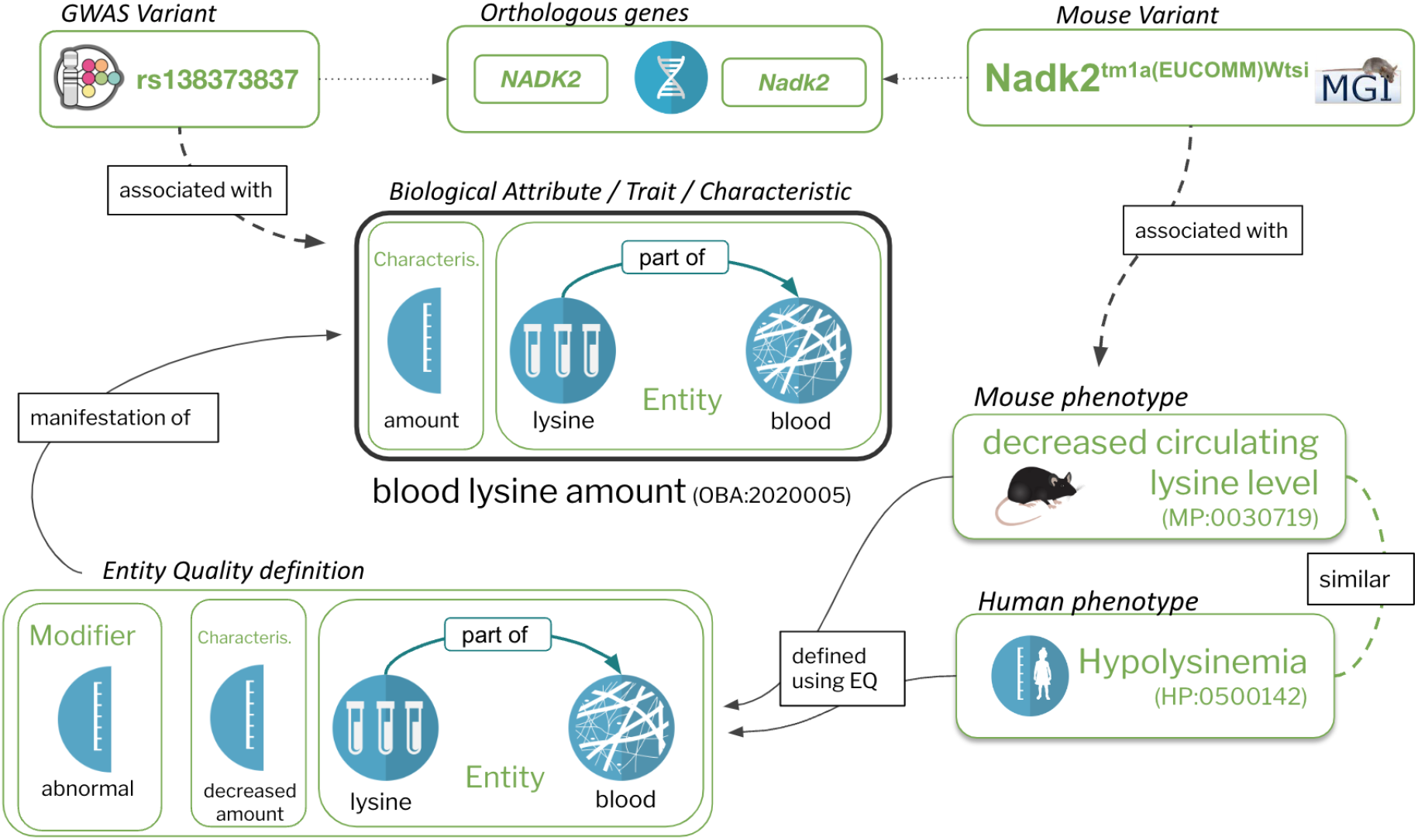
The Entity-Quality model enables composing biological attributes in a way that is compatible with the logical definitions of widely used ontologies such as the MP and HPO which are used to document phenotypes associated with diseases or genes. On the right is a specific example of a human phenotype term, “Hypolysinemia” (HP:0500142), which means a lower than normal amount of lysine in the blood. The EQ (phenotypic effect) on the left is not only used to logically define Hypolysinemia, but also the mouse phenotype “decreased circulating lysine level” (MP:0030719). This ensures that an automated reasoner can compute the appropriate relationship between the two (in this case equivalence), as well as to the specific biological attribute they are concrete manifestations of (“blood lysine amount”). Representing phenotype and phenotypic attributes this way enables the grouping of quantitative variant data (e.g. GWAS) and qualitative variant data (e.g. MGI).

OBA is a standardised, representational framework for observable attributes that are characteristics of organisms, or parts of organisms. For example, the attribute “trochanter size” (OBA:0002360) is a characteristic of the anatomical entity “trochanter” (UBERON:0000980); and “blood glucose amount” (OBA:VT0000188) is a characteristic of glucose (CHEBI:17234) in the blood (UBERON:0000178). This way of defining attributes, called the Entity-Quality (EQ) pattern, is used by many biomedical ontologies, including the Plant Trait Ontology (TO^19^) for defining attributes of plants such as “petal length” (TO:0002605), the Environment Ontology (ENVO)^20^ for defining attributes of environmental materials, such as “soil pH” (ENVO:09200010), and the Human Phenotype Ontology (HPO) for defining phenotypic abnormalities such as “Abnormal telomere morphology” (HP:0031412). The same EQ pattern has also been employed in data annotation using a post-compositional approach—combining an entity term and a quality term within an annotation, rather than creating a separately defined trait term—to describe both phenotypic abnormalities (e.g., in zebrafish^21^) as well as natural evolutionary variation in the Phenoscape Knowledgebase^22,23^. The initial design of OBA was significantly inspired by work from the creators of the Plant Trait Ontology.

The majority of attributes in OBA are logically defined using the Web Ontology Language (OWL). These logical definitions use terms from relevant reference ontologies, such as Uberon^24^ or ChEBI^25^. With the exception of a small number of high-level concepts, most of the classification in OBA is automatically computed on the basis of the classifications of the various reference ontologies, using an automated reasoner. The advantage of this approach is twofold: firstly, we do not have to manually classify any concepts, which drops the cost of curating the classification significantly while increasing its completeness. Secondly, the numerous links to reference ontologies can be exploited for a wide variety of applications, including querying (e.g., select all data where the morphology of a part of the renal system is affected), knowledge graph integration (e.g., automatic linking to phenotypic abnormalities from widely used ontologies such as HPO or MP), and knowledge inference (e.g. inferring missing data from logical implications^23^). A rich logical axiomatisation based on design patterns is necessary to ensure interoperability with existing phenotype ontologies and other data types, such as anatomical, chemical and biological pathway data. Existing ontologies such as the Vertebrate Trait ontology (VT)^26^ and the Experimental Factor Ontology (EFO)^27^ are widely used to annotate traits, but do not contain such axiomatisation.

In this paper, we introduce OBA, an ontology and logical framework for representing biological attributes. We show how we use the Entity-Quality modelling framework to automatically classify attributes reliably using reference ontologies. Additionally, we demonstrate how OBA can be used to automatically integrate data from other widely used phenotype ontologies, thereby breaking silos.

## Methods

### Logical Framework

As ontologies grow in size, they become increasingly hard to maintain. Phenotype (and trait) ontologies are inherently polyhierarchical, as they combine a variety of interwoven external classifications, such as attributes and biological entities. This makes it hard to ensure that the classification is complete by manual curation (no subclass axioms are missing), and that the existing classification is consistent with other ontologies (for example, “head size” should not be a parent of “eye size”). Instead of relying on manual classification of biological attributes in OBA, we use logical definitions and automated reasoning to compute the hierarchical classification. OBA is represented in the Web Ontology Language (OWL), a knowledge representation formalism based on Description Logics, a fragment of First Order Logic. It is fully aligned with the Core Ontology for Biology and Biomedicine (COB^28^) because all concepts in OBA are, implicitly, children of “characteristic” (PATO:0000001), which itself is part of COB. However, we currently do not import COB directly (though this is planned as future work).

The Entity-Quality (EQ) pattern^29^,30 is widely used for representing traits and phenotypes in ontologies such as the Human^18^, Mammalian^17^ and Xenopus^31^ phenotype ontologies. There are a number of variants of this pattern, but at its core, a phenotypic quality (Q, which is currently more frequently referred to as a “characteristic” rather than “quality”) such as “height”, “mass” or “amount”, usually from the Phenotype And Trait Ontology (PATO)^32^, is combined with an entity (E), such as an anatomical or chemical entity, to form the concept of a “biological attribute”, sometimes referred to as a “trait” (see Figure 1). For example, “lysine in blood amount” (OBA:2020005) is composed of the PATO class of “amount” (PATO:0000070), lysine (CHEBI:25094) and blood (UBERON:0000178) (see Figure 1). PATO defines basic categories of phenotypic qualities (attributes or characteristics) and it can be used for quantitative trait or Mendelian phenotype annotation^33^. PATO is species-neutral in its scope but does not provide relationships to the biological entities whose phenotypic qualities it is meant to describe^30^. Using EQ logical definitions in OWL enables us to use automated reasoners to automatically classify our traits: if, for example, lysine is an “amino acid” according to ChEBI, there is no need to remember to manually classify “lysine in blood amount” under “amino acid in blood amount” - the reasoner will do this for us based on the classification in ChEBI. A second feature of such axiomatisation is that it can be used for powerful logical querying using OWL DL Queries ^34^ and SPARQL^35^. This enables us to group data in ways that are not easily available in traditional databases. For example, it allows us to query for data related to morphology of a tissue that is considered part of the cardiovascular system - even if no such term exists in OBA. A query capturing this can be found in the supplemental materials (S2, supplemental materials).

Ontologies of phenotypic abnormalities such as HPO, MP, XPO^31^ and ZP make extensive use of the EQ pattern, but are primarily used to capture phenotypic effects compared to some reference (usually “wild type”) rather than unqualified biological attributes as in OBA. For example, “decreased circulating lysine level” (MP:0030718) in the Mammalian Phenotype Ontology is defined as “abnormal(ly)” (PATO:0000460) “decreased amount” (PATO:0000470) of “lysine” in the “blood” (Figure 1). Since both the biological attributes in OBA and the phenotypic effects in MP are represented using the Web Ontology Language (OWL), we can use an automated reasoner, such as ELK^36^, to automatically compute links between the two. Other examples of links between OBA attributes and phenotypic effects: head circumference (OBA:VT0000047) has “Decreased head circumference” (HP:0040195), “Microcephaly” (HP:0000252) and “Progressive microcephaly” (HP:0000253) as manifestations; “brain ventricle size” (OBA:0002294) has manifestation “Ventriculomegaly” (HP:0002119).

### Template-based ontology curation with DOS-DP

Ontologies, especially those with logically rich axiomatisation, enable powerful services such as automated reasoning, classification and logical querying, but logical modelling is difficult^37^ and appropriate expertise is scarce. A popular approach to deal with this problem is to use design patterns and templating systems for logical axioms^38^. This allows for decoupling the curation of reference terms used for logical definitions from their exact axiomatic pattern. The central idea is to employ a small number of axiom templates (which implement design patterns such as the EQ model described above) that can be created and maintained by logic experts, and have content curators focus on the selection of appropriate filler terms (e.g. terms from Uberon to define anatomical attributes). There are a number of available approaches, but many OBO Foundry ontologies use Dead Simple Ontology Design Patterns (DOS-DP^38^), a system that allows us to capture the logical model in a specific YAML file (design pattern), which is separate from curation of the actual biological attributes. For example, the “entity attribute” template (see Figure 2), the most basic of all OBA design patterns, has two filler terms, the entity (e.g. a chemical, or an anatomical entity) and the attribute (a characteristic from PATO, such as “amount”) and defines how a new biological attribute in OBA following that pattern should be converted to OWL (among other aspects, it describes how the logical equivalent class definition should be instantiated). Curators simply add a row to a spreadsheet with the OBA identifier and the two fillers. The identifier scheme used for new OBA terms corresponds to the standard OBO recommendation^39^. All identifiers are represented as an OBO PURL, starting with the http://purl.obolibrary.org/obo/OBA_URI_prefix, followed by an numeric identifier. Some identifiers are prefixed with the literal “VT” to indicate that they are sourced from VT. A specialised toolkit (DOS-DP tools^40^) then translates the spreadsheet into OWL axioms using the template file.

**Figure 2:**
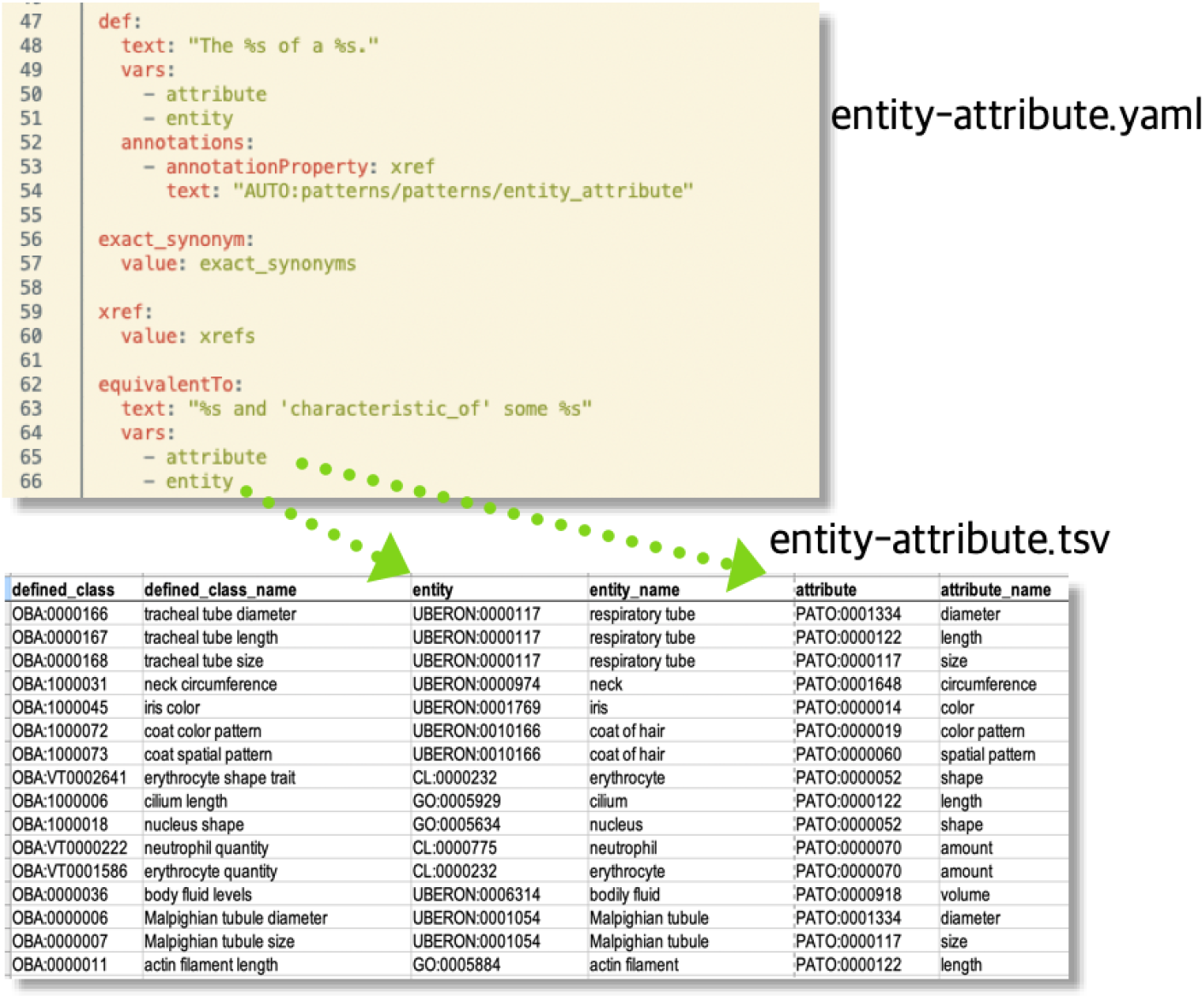
DOS-DP template example. The fillers declared in the template above (attribute, entity) are mapped to the respective column names in the TSV file below. A specialised tool reads both files and generates the axioms specified by the template file.

OBA currently uses ten DOS-DP term templates for different trait patterns; see Table S1 (supplemental materials). These were selected because they cover the majority of anatomical, chemical level and cellular attributes which are central for the integration of genomics data. By far the majority of biological (especially anatomical) attribute terms in OBA can be represented using a basic entity-attribute pattern (e.g. “head size”). All templates can also be found online^41^. In addition to ensuring a consistent axiomatisation of the ontology across thousands of terms (a general advantage of template systems, not just DOS-DP), one major advantage of using DOS-DPs as a framework for managing OWL ontologies is their generative capabilities. Not only can we dynamically generate labels, definitions and synonyms based on the filler terms provided, but we can also add contextual axioms which can be exploited for automated reasoning. In OBA, for example, we generate General Concept Inclusion (GCI) axioms which define how attributes are related mereologically (e.g. “ulna size” part of “forelimb skeleton size”). These axioms are defined as part of the DOS-DP design patterns.

### Automated mapping pipeline for external sources

One of the central use cases for OBA is to provide additional structure to other generally weakly axiomatised ontologies, mainly the Vertebrate Trait Ontology (VT) and the Experimental Factor Ontology (EFO). To synchronise these vocabularies with OBA, we execute the following workflow:

1. Match: link external terms to OBA terms if they exist
2. Sync: identify external terms that do not exist in OBA, decompose them into their logical components (reference entities) and curate them as instances in our DOS-DP template pipeline
3. Compile: compile the new terms into OWL and integrate them into the ontology

We have built a custom pipeline^42^ that supports our curators in steps 1 and 2. To that end, we implemented a matching process that works on the ontology labels and exact synonyms. After applying a series of normalisation steps (including the removal of stop words like “measurement” or “trait”), if a direct match between an external term and an OBA term can be identified, we present it as a candidate match to a curator. The curator just has to review and approve or reject the match. For step 2, we sequentially match all our reference ontologies (ChEBI, Uberon, PRO, GO, PATO) to the external term. For example, if an external term “lysine measurement” contains the term “lysine”, we record that as a potential match for the “entity” column in the “entity-attribute” DOS-DP pattern (Figure 2). Thus our curators are presented with a set of potential EQ-decompositions, which they proceed to either accept or reject. Mappings to external ontologies, as generated in steps 1 and 2, are documented using the Simple Standard for Sharing Ontological Mappings (SSSOM)^43^ and shared as part of the OBA GitHub repository (https://github.com/obophenotype/bio-attribute-ontology). Note that in contrast to other synchronisation workflows such as those used by Uberon or the Mondo disease ontology ^44^, we do not import any curated information from external ontologies (synonyms, definitions, etc.) but rely entirely on automated templated processes.

### OBA life cycle management

OBA has been a member of the OBO Foundry^45^ for more than seven years and has a team of 6 regular contributors. It is managed by members of the European Bioinformatics Institute (EBI) and the Monarch Initiative^46^ using modern ontology workflows and curation practices. To manage our releases, quality control and external dependencies we use the Ontology Development Kit (ODK^47^, version 1.3.2). The ODK provides mechanisms to version and publish OBA releases in a variety of serialisations (JSON, RDF/XML, OBO) and release file variants according to OBO Foundry practices, relying largely on the ROBOT tool^48^. It fully supports DOS-DP workflows which ensures a seamless integration of mostly TSV based curation into the general ontology life cycle. For example, terms that are used as fillers during the decomposition of biological attributes are automatically imported from their respective external ontologies. The ODK is also used for continuous integration testing. Whenever one of our curators makes a pull request on GitHub with changes to OBA, we automatically execute the DOS-DP pipeline, followed by a number of strict quality control checks. For example, these checks ensure that all terms added fall under the “biological attribute” root term, are unique (no other equivalent attribute exists) and are logically consistent. Lastly, the ODK imports relevant terms and axioms from our reference ontologies (e.g. Uberon, ChEBI, RO), which ensures that OBA is fully consistent with their axiomatisation. To ensure consistency, we use the ELK reasoner, which is suitable for OWL 2 EL ontologies (see Results). OBA publishes a new version every 2-3 months, using the GitHub releases mechanism for versioning and dissemination.

## Results

The Ontology of Biological Attributes (OBA) is published under the CC0-1.0 licence (public domain) and is in its 17th release (21 December 2022)^49^ at the time of writing this paper. The ontology is expressed using the OWL 2 EL profile of the Web Ontology Language (OWL^50^). Note that some imports use higher expressivity axioms (beyond OWL 2 EL), which means that there are corner cases where using an OWL 2 EL reasoner such as ELK^36^ may be incomplete. (Note: all elements required by the Minimum Information for Reporting of an Ontology (MIRO) guidelines^51^ are reported here.)

OBA defines 7807 biological attributes, most of which have logical equivalence axioms (full logical definition).

OBA attributes have, on average, 1.54 parents (indicating a high degree of polyhierarchy) and one or more associated synonyms. Most attributes (69.6%) are anatomical, 12.4% are attributes of biological processes and 9.8% are cellular attributes (see Figure 3). Anatomical attributes are defined with terms from the Uberon anatomy ontology. In total there are 302 CHEBI, 346 CL, 708 GO, 40 MONDO, 109 NBO, 139 PATO, 69 PRO, 4 SO and 2469 UBERON terms referred to by OBA ids. The latest version of OBA is available under the persistent URL http://purl.obolibrary.org/obo/oba.owl. The version referred to as part of this paper can be accessed by the versioned persistent URL http://purl.obolibrary.org/obo/oba/releases/2022-12-21/oba.owl.

**Figure 3:**
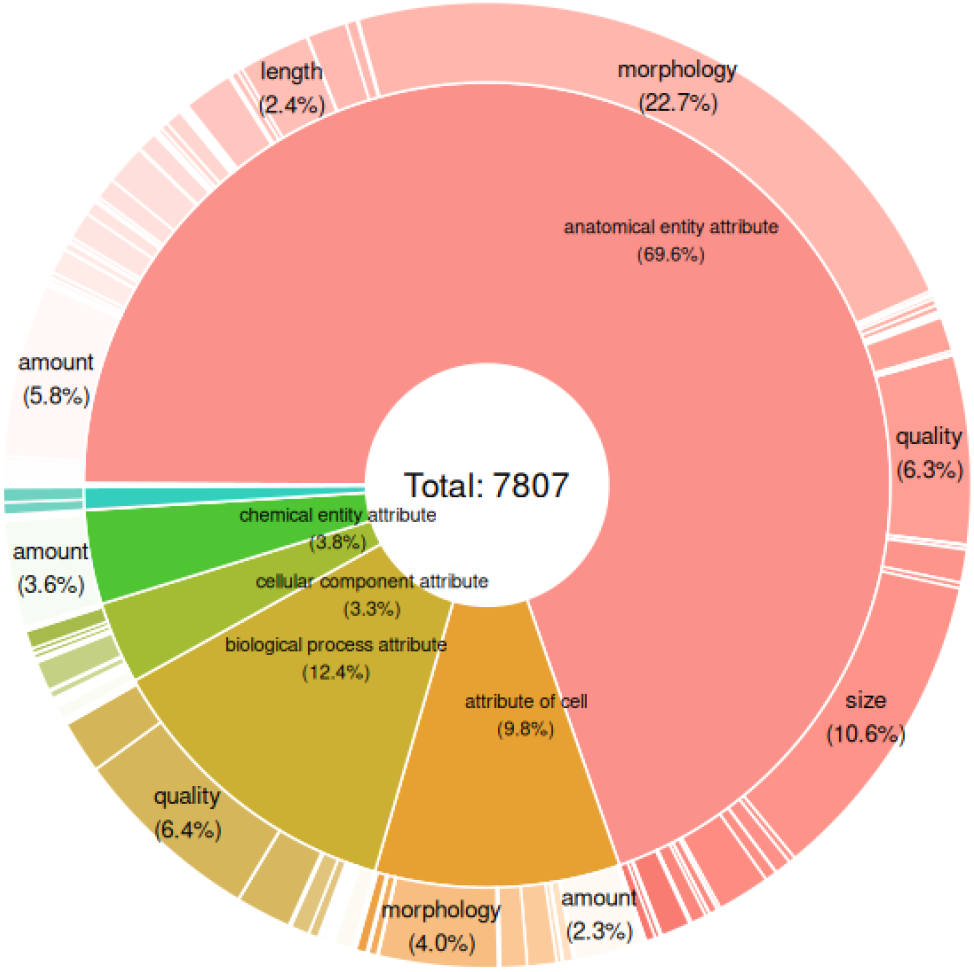
Distribution of OBA attributes across categories and qualities.

If an entity is obsoleted, the label is changed to have “obsolete” at the beginning, the metadata field “deprecated” is true, and all logical axioms are removed. When the entity is completely redefined, the metadata field “term replaced by” is added to indicate the substitute term.

The four main relationships used in OBA are “part of” (BFO:0000050) primarily to denote mereological links between anatomical entities, “characteristic of” (RO:0000052) to link a characteristic (e.g. “morphology”) to a biological entity (e.g. “heart”), “characteristic of part of” (RO:0002314) to link a characteristic to a biological entity and all of its parts (e.g. we can use “characteristic of part of” to define a trait that applies to all parts of the cardiovascular system), and “subclass of” (rdfs:subClassOf) to classify biological attributes. In addition, the relationship has_role (RO:0000087) is used in cases where the definition of a biological attribute requires a reference to a chemical role, such as “serum metabolite” in “serum metabolite amount” (OBA:2050092).

### OBA coverage of biological attributes relevant for cross-species data integration

The template-based curation workflow (Section “Template-based ontology curation with DOS-DP”) makes the process of adding new attributes highly scalable, as we do not need to worry about logical modelling. In the following we show (as an example) how to rapidly curate relevant biological attributes in OBA to cover the needs of the International Mouse Phenotyping Consortium (IMPC) database which captures information that includes the effect of gene knockouts on phenotype^7^. IMPC uses 1102 phenotypic abnormality terms from the Mammalian Phenotype Ontology (MP). To ensure that we capture the relevant attribute terms for these, we first extract the fillers for the EQ logical definitions from MP using DOS-DP tools^52^, and then transform the fillers into the respective OBA pattern. 532 (∼50%) of the IMPC phenotypes could be matched this way. A curator then manually assigns appropriate PATO characteristics (e.g, “amount” in cases where the quality of the phenotypic abnormality was “increased amount”). This process resulted in a total of 179 new trait terms added to OBA.

The Mouse Phenome Database (MPD)^14^ enables the integration of genomic and phenomic data by providing access to primary experimental data, well-documented data collection protocols and analysis tools. OBA terms currently cover 80.1% (5066 of 6325) of trait measures annotated in MPD via mappings to VT. We estimate that we will cover most of the remaining 20% by the end of 2023.

Identifying mouse traits reflective of human disease is critical to prioritise preclinical models of disease and aspects of complex disease. Prior to the development of OBA, researchers hoping to retrieve mouse trait measures reflective of human disease characteristics had to know specifically which mouse traits were associated with each disease, searching trait by trait to find a complete set. A workaround approach involves retrieval of disease terms (DOID) to vertebrate traits (VT) using a gene-centric mapping performed by retrieving Alliance of Genome Resources (AGR)^4^ annotated genes associated with each DOID and identifying Mammalian Phenotype (MP) terms to which their phenotypic alleles were annotated in Mouse Genome Database (MGD)^5^. These MP terms were used to retrieve mouse trait measures from the Mouse Phenome Database (MPD) and the VT terms to which the traits were also annotated. In an effort to validate the utility of OBA to retrieve relevant mouse data, we compared this gene-centric approach to the OBA’s semantic mappings of disease terms from the Disease Ontology (DO)^53^ to vertebrate traits (VT). In total, OBA mapped 3033 of 3455 (87.8%) disease terms to VT terms with mouse trait measures in MPD. From the two approaches combined, we identified 4910 disease terms that had associations to VT terms. For 1348 disease terms, at least one VT term was mapped to the disease using each approach and 558 (41.4%) were associated with at least one shared VT term. In these cases, the mean overlap of VT terms per disease was 26.6% for OBA and 16.7% for the gene-centric approach.

### Using OBA to group phenotypic abnormalities

To determine how well existing ontologies of phenotypic abnormality aggregate under OBA, we count the total number terms falling under any OBA class, and the total number of links between a phenotype ontology and OBA. For an accurate edge count, we follow the Ubergraph approach, which essentially converts an ontology to a knowledge graph with nodes and edges instead of axioms. We (1) merge the phenotype ontologies with OBA, then (2) materialise the relationships necessary for connecting OBA biological attributes with phenotypic abnormalities using a regular OWL 2 reasoner (ELK). Next, we (3) convert the resulting ontology to a knowledge graph using “relation graph”^54^. Lastly, (4) we extract the OBA mappings from the knowledge graph using the SSSOM toolkit^43^. The results can be found in the table below. It is important to understand that no particular effort was made to cover all phenotype classes - as described in the section above, coverage can be rapidly increased by reusing the logical definitions. This experiment only illustrates the breadth of integration, not its depth: it includes classes that only link to very general attributes like “morphology anatomical entity”.

### Alignment with other trait ontologies

In contrast to the alignment with ontologies of phenotypic abnormalities as described in the previous section, alignment with most other trait vocabularies has to be performed using a semi-automated approach based on automated matching (see Section on “Automated mapping” above) and manual curation. To date, we have curated 2314 mappings to VT (version 12.5) and 150 mappings to EFO (version 3.14.0). 2,332 terms in OBA (which can be recognised by their ID, i.e. OBA:VT123 instead of OBA:123) have been derived from the VT ontology, i.e. OBA terms decompose and generalise them using the EQ pattern.

A handful of other ontologies of attributes use the same EQ system to define attribute classes. For example, the Plant Trait Ontology (TO) has 1144 classes that classify under OBA attributes, and the Plant Phenology Ontology has 60 such classes.

### How to access OBA and how to contribute to it

The EMBL-EBI Ontology Lookup Service (OLS)^55^ and Ontobee^56^ are platforms from which one can find or browse OBA terms manually. There is also an OBA GitHub repository for those who wish to contribute to OBA, view documentation or download public releases and source files.

Users can explore OBA by entering free text into the search box on OLS or by using unique, permanent OBA identifiers. It is also possible to browse terms in the ontology hierarchy tree view or the interactive graph layout which displays colour-coded term relations. Search results return OBA terms, textual and logical definitions in addition to terms dynamically imported from other ontologies. Users can also query for OBA terms using the linked ontology server, Ontobee (Table 2). A complete list of terms can be downloaded in “.xlxs” or “.tsv” formats from Ontobee’s home page. The OBA “.obo” or “.owl” ontology files can be viewed in an ontology editor such as Protégé, where users can browse terms and construct DL queries^57^.

**Table 1:** Current degree of integration between OBA and existing phenotype ontologies

| Ontology | # Links to OBA | # Classes under OBA |
| --- | --- | --- |
| HPO | 217,474 | 16,544 |
| MP | 187,405 | 13,620 |
| ZP | 117,023 | 39,373 |
| XPO | 38,567 | 20,340 |

**Table 2:** Different ways to access OBA

| Link | Name | Note |
| --- | --- | --- |
| https://www.ebi.ac.uk/ols/ontologies/oba | EMBL-EBI Ontology Lookup Service (OLS) | Find or browse OBA terms manually. |
| https://github.com/obophenotype/bio-attribute-ontology | OBA GitHub repository | To contribute to OBA, read documentation, or download OBA public releases and source files. |
| https://www.ebi.ac.uk/ols/docs/api | OLS API | Query OBA terms programmatically. |
| https://ubergraph.apps.renci.org/sparql | ubergraph SPARQL endpoint | Query OBA terms programmatically. |
| https://api.triplydb.com/s/DwuipbH9o | Example query using ubergraph SPARQL endpoint | Query OBA terms programmatically. |
| https://ontobee.org/ontology/OBA | Ontobee | Query OBA terms manually or programmatically. |
| https://github.com/obophenotype/bio-attribute-ontology/issues | OBA issue tracker | Contribute bug reports or suggestions to OBA. |
| https://github.com/INCATools/ontology-access-kit/blob/main/notebooks/Monarch/OBA-Tutorial.ipynb | Accessing OBA using the Ontology Access Kit (OAK) | We provide a Jupyter notebook showing examples of querying of OBA using OAK. |

OBA welcomes contributions or suggestions for improvements from the research community. Contributions, suggestions or bug reports can be initiated via the OBA issue tracker on GitHub (Table 2).

### Programmatic access to OBA

OBA is distributed in RDF/OWL, OBO Format, and OBO Graph JSON format, so any programming library that is capable of reading these formats can be used to explore OBA. For data science use cases we recommend the use of the Ontology Access Kit (OAK)^58^, which provides both Python bindings and a command line interface. Additionally, there are several ways to query OBA terms using public APIs: the OLS API, Ubergraph^54^ and Ontobee SPARQL endpoints (Table 2).

### Use cases

OBA is used across a wide range of biological domains and processes, including genomics and drug discovery. In this section, we list some examples of its use.

The Gene Ontology^59^ is using OBA for axiomatising their “regulation of characteristic” branch (∼100 terms), which describes biological processes that qualitatively or quantitatively modulate a biological attribute. For example, biological processes “regulation of lysosomal lumen pH” (GO:0035751) and “lysosomal lumen pH elevation” both regulate the biological attribute (trait) “lysosomal lumen pH” (OBA:0000091). OBA trait terms imported into EFO can facilitate computational drug target identification via the Open Targets Platform^60^. For example, OBA, in tandem with other ontologies, has proved useful for computational drug target identification in a study of drug-induced adverse events in animal models ^61^.

Another important OBA use case is the online community resource Functional Trait Resource for Environmental Studies (FuTRES)^62^. It contains an application ontology, FuTRES Ontology of Vertebrate Traits (FOVT) (https://obofoundry.org/ontology/fovt.html), developed to standardise measurable trait terms in vertebrates. The FOVT currently has 390 trait terms (https://futres-data-interface.netlify.app/), 65 of which are from OBA and 325 of which will be eventually incorporated into OBA^62^. By standardising terms, researchers spend less time wrangling data as the harmonised terms enable interoperable data. FOVT follows and helps develop patterns developed by OBA. Using patterns helps eliminate human errors and makes for easier on-boarding of new ontology curators. FuTRES also takes advantage of the annotation properties in OBA, such as annotations for taxon- or field-specific trait terms, to increase findability by researchers of trait terms.

OBA terms are also used in the fields of agriculture, nutrition, zoology and biodiversity. AgBioData member databases take advantage of the species-neutral nature of OBA terms to integrate agriculturally important animal and plant traits with genomics and genetics data^63^. The Compositional Dietary Nutrition Ontology (CDNO) uses OBA to link nutritional components found in food to their human dietary roles which include traits. This allows the integration of nutritional components with associated traits, for example, bone strength (OBA:VT0001542)^64^. OBA trait classes have been used for the annotation of domestic guinea pig electrophysiology data^65^. The Encyclopedia of Life (EOL) TraitBank takes advantage of the well-axiomatised OBA terms to infer traits in biodiversity data and to improve their search functionality ^66,67^. For example, a user looking for a body size measurement would not have to do separate searches for all the different ways body size is measured in different taxonomic groups, e.g., body length, snout vent length, fork length, etc. The semantic features of OBA can contribute to improved named entity recognition performance when incorporated in a natural language processing (NLP) framework for biodiversity literature curation^68^. Additionally, OBA can be used to link traits and phenotypes to environments^69^. This is of particular interest in the crop science community, where researchers are working to identify specific regions of the genome that control complex traits, such as drought resistance.

Having a rich set of links between biological attributes (traits) and phenotypic abnormalities also enables a wide range of applications. For example, we can use these links to group data across species at a high level. Many databases such as the Mouse Phenome Database (MPD) have to deal with the challenge of grouping trait data from a variety of studies for meta-analyses, e.g., all trait data associated with hypertension should be grouped. Similarly, the hierarchical structure allows for a broader search in FuTRES, where a user can query for “humerus length” and have all the ways humerus length is measured with measurement values returned, rather than having to do a separate search for each measurement method to retrieve data.

OBA is a component of the successor of the Unified Phenotype Ontology^70^, uPheno 2^71^. uPheno 2 is used by the Monarch Initiative to integrate gene-to-phenotype data across species. The integration of OBA allows grouping of phenotype data across traits without concern for the specific manifestation (e.g. “blood glucose level” instead of “abnormally increased blood glucose level” or “limb morphology” instead of “short limb”, etc.).

### Limitations of the approach

#### Identifier uniqueness

One of the key OBO Foundry principles is class uniqueness: a single term, such as “amount of lysine in the blood”, should not exist in multiple ontologies. While the synchronisation with EFO is unproblematic (EFO is not an official OBO Foundry reference ontology, and measurement terms are conceptually disjoint from trait or attribute terms), the synchronisation with VT may raise some questions. While the class uniqueness principle is absolutely central to reference ontologies such as PATO, ChEBI, GO and PRO, it is very complicated to maintain in a cross-species context. The prevalent practice is to have one species-independent vocabulary (Uberon, uPheno, and now OBA) whose goal it is to integrate species-specific ontologies (MA, VT, XAO, ZFA, FOVT, etc.). Furthermore, species-specific ontologies are typically maintained as taxonomical structures (owl:subClassOf hierarchies with little additional axiomatisation) which means that they lack the strong logical foundation that integrator ontologies provide.

#### Need for manual mapping curation

The integration of GWAS data with data from a more qualitative phenotyping pipeline relies to a large extent on our mappings between OBA and EFO, and VT, which is an ongoing process. Due to their lack of (logical) formalisation, alignment is largely manual, but the comparatively small sizes of the relevant branches in EFO and VT makes it feasible to curate mappings semi-automatically using automated matchers and manual curation, as described in the Section on “Automated Mapping” (Methods). The rapid improvement in Large Language Models and other NLP techniques may be able to speed up this process in the future.

## Discussion and future work

The primary objective of OBA is to break silos across data types related to characteristics (e.g. “amount” or “mass”), traits or biological attributes (e.g. “amount of lysine in blood”), phenotypic abnormalities (e.g. “Hypolysinemia”) and biological entities/processes (e.g. “blood”, “lysine”, “mitosis” or “cardiovascular system”). Due to its rich logical definitions, OBA naturally integrates well with data focused on links to anatomy (such as gene expression data), chemical entities, cellular components, cell type, cell types, biological process and more. This allows, for example, the integration of anatomy focused data (such as gene expression and single cell expression data) with trait-level data which is already a significant improvement over the status quo. Existing vocabularies to capture biological attributes, such as VT and EFO, do not (aside from the provision of simple cross-references) systematically bridge the gap between PATO characteristics, reference ontologies (e.g. anatomy, chemical), and phenotype ontologies.

Phenotype ontologies such as MP and HPO that define phenotypic abnormalities have been used for over a decade in the biomedical domain for clinical and model organism phenotyping. Due to the widespread use of the EQ design pattern (see the “Logical Framework” Section), we can classify phenotypic abnormalities under their respective biological attributes (Section “Using OBA to group phenotypic abnormalities”). Public endpoints such as Ubergraph (see Section “Programmatic access to OBA”) demonstrate how hundreds of thousands of links between biological attributes and phenotypic abnormalities can be inferred automatically without a human in the loop. Furthermore, the integration of OBA into uPheno 2 allows to easily group phenotypic effects across biological attributes, which opens up powerful possibilities for search and grouping of annotations (Section “Use cases”).

Many polygenic, quantitative, and GWAS traits are not in scope for the Mendelian phenotype focussed ontologies. There are ontologies that focus on or include quantitative and measurable trait terms, such as VT, EFO and the Clinical Measurement Ontology (CMO)^72^,73. EFO curators, for example, maintain a branch in the ontology for “measurement” terms that are used in annotation by the GWAS Catalog^13^. “Measurement” terms, such as “urinary sodium measurement” (EFO:0021522), have a broad applicability in annotation. They can be used to annotate experiments independent of any conclusion about the results and outside of any context where conclusions might be made about biological traits. The GWAS Catalog uses these terms to record something more specific - an association between the presence of an allele and some effect on the measured value of a trait. Mapping a GWAS annotation with a “measurement” term to an OBA term, such as “urine sodium amount” (OBA:VT0006274) enables recording this explicitly, and has the advantage that the terms can be integrated directly with widely used phenotype ontologies, e.g. “Hypernatriuria” (HP:0012605). Measurement terms are still useful as they can record one of many assay methods for measuring a specific trait. For example, Body Mass Index is a useful, if sometimes limited, proxy measurement of body fat levels. Using a BMI measurement term to annotate GWAS variants can record this useful information, mapping this to a trait term for body fat levels then allows this to be integrated with related traits and phenotypes. Specialised ontologies that capture the measurement method exist, for example the Ontology of Biomedical Investigation (OBI)^25^ or the Biological Collections Ontology (BCO)^74^.

The integration of quantitative trait data (such as GWAS or QTL) with outcomes from clinical and research organism phenotyping activities is one of the most promising applications of OBA. For example, the deep integration between OBA and HPO will facilitate the use of gene-phenotype associations derived from GWAS studies in variant prioritisation software such as Exomiser^75^, which is used for clinical diagnostics. This has the potential to significantly extend the existing sources of gene-phenotype data from annotations of Mendelian disease resources such as OMIM and Orphanet as well as model organism resources such as MGI^5^, IMPC^7^ and ZFIN^76^.

As the space of biological attributes / traits is very large, any curation of new terms must be highly scalable. The cost of manually classifying biological attributes and phenotypes is high (leading to incomplete classifications). To demonstrate how defining new biological attributes can largely be automated, we rapidly aligned more than 500 terms from the Mammalian Phenotype Ontology with OBA (Section “OBA coverage of biological attributes relevant for cross-species data integration”) by repurposing logical definitions used and focussing on the curation of the specific phenotypic characteristic (e.g. “amount” instead of “increased amount”). Using logical definitions for automated reasoning and templates for scalable curation enables rapid development of terms.

OBA can facilitate the interpretation of trait and phenotypic findings in clinical laboratory test results, many of which are annotated with Logical Observation Identifier Names and Codes (LOINC)^77^. As part of future work, we will bridge OBA to the LOINC database via the CompLOINC project (https://github.com/loinc/comp-loinc), which decomposes the (heavily pre-coordinated) LOINC classification into an OWL ontology with is-a hierarchies for each of the 6 LOINC Part Types (Component, System, Method, Property, Time and Scalar). This OWL formalisation of LOINC allows logical reasoning, subsumption querying by Part Type, and has the potential to provide an extensive bridge between the LOINC-dominated clinical laboratory domain and the phenotype ontology world that dominates in the area of genomics. Rather than matching LOINC codes on the level of the highly variable LOINC labels, OBA terms can be matched much more easily to LOINC Part terms (e.g. chemical entities to ChEBI, anatomical entities to Uberon). This will result in a much-needed bridge between the clinical laboratory domain and biological research and genomics.

Also as future work, we seek to integrate OBA with disease ontology terms (which are also widely used, for example, in GWAS) through phenotypic features of diseases and common links to Uberon. For example, “familial juvenile hyperuricemic nephropathy” (MONDO:0000608) is linked to “Hyperuricemia” (HP:0002149) which is logically defined as “increased amount of uric acid in the blood”. It therefore is automatically classified under “blood uric acid levels” (OBA:VT0010302). This gives us a natural bridge from diseases to biological attributes which provides another layer of integration. A second level of integration that has yet to be explored is to exploit the numerous “anatomical site” relations provided by disease ontologies such as Mondo - these are already integrated with OBA through the use of a common reference ontology (Uberon), but biological attribute terms could easily be generated based on the Uberon reference for more thorough logical integration. Prior to the development of OBA, researchers hoping to retrieve mouse trait measures reflective of human disease characteristics had to know specifically which mouse traits were associated with each disease, searching trait by trait to find a complete set. Using OBA mappings, mouse trait measures in MPD, for example, are readily annotated to 3033 disease ontology terms with more than 50% coverage across all MPD trait measures, allowing researchers a simple means of retrieving all disease associated trait data using a single, intuitive disease-centric query. These data can then be used to identify preclinical mouse models collectively extreme across a set of disease-related traits.

### Related work

The Vertebrate Trait Ontology (VT)^26^ is a cross-species, unified trait vocabulary used for the annotation of terms in vertebrates, with a goal of standardising vocabularies to enable interoperable data and cross-study comparisons. It was created based on the structure of the Mammalian Phenotype Ontology (MP), where references to abnormalities were removed and a skeletal set of neutral trait terms was maintained after a manual review. It is therefore a phenotype-neutral ontology, which, similar to OBA, describes traits that do not indicate an abnormal state or process or express any phenotypic variation. It is used by the Mouse Phenome Database (MPD^14^) for the annotations of all strain measurements. The Mouse Genome Informatics (MGI), Rat Genome Database and Animal QTL Database use VT terms for the annotation of Quantitative Trait Loci (QTLs). VT cross-references to other ontologies, such as the Gene Ontology (GO), by transitively connecting terms using the “is_a*”* relationship. Unlike OBA, VT terms are not constructed using logical axioms, and there are no logical links to other ontologies. VT uses weak non-logical cross-references to GO and MP to indicate that a link exists, but these links are sparse and cannot be used for automated reasoning (less than 20% of VT terms have such links, compared to 100% of OBA, which are logical and therefore more meaningful).

The Zebrafish Information Network (ZFIN) is the central knowledgebase for the species *Danio rerio*, providing genotype, phenotype and disease models data. ZFIN has curated more than 52,000 phenotype annotation statements that are constructed using the Entity-Quality (EQ) syntax, employing terms from the Zebrafish Anatomical Ontology (ZFA), PATO, GO and ChEBI ontologies. It is a member of the Alliance of Genome Resources, as well as a core member of the uPheno initiative, which works towards the large-scale reconciliation of divergent logical definitions across species^70^.

The Plant Trait Ontology (TO)^19^ is a Planteome database reference ontology that describes phenotypic traits in plants, linking ontologies and facilitating cross-species studies. It is species-neutral; each trait is a distinguishable feature, characteristic, quality, or phenotypic feature of a developing or mature plant. Plant traits, which are defined in TO as measurable characteristics, scale from molecular entities to plant cells and types to whole populations. TO was first created to describe QTL traits and its current hierarchical structure describes nine upper-level plant trait classes. Many TO terms also follow the EQ pattern, drawing entities from ontologies like the Plant Ontology (PO), GO and ChEBI and quality terms from PATO to provide pre-composed descriptions of terms and logically connect TO to other ontologies.

The Animal Trait Ontology (ATO)^78^ was created in 2008 as an effort to form a central, standardised repository of controlled, phenotypic trait terms for three domesticated farm animal species. As in many trait ontologies, traits are defined by ATO as being measurable, while phenotypes are expressed as scalar with directionality. ATO uses the “is_a” and “part_of” axioms to describe relationships between terms. ATO was later expanded and is now referred to as the Animal Trait Ontology for Livestock (ATOL), an ontology of traits defining phenotypes described in the Environment Ontology for Livestock (EOL). Its top-class trait term “animal trait of livestock” has seven main branches and now includes species ranging from ruminants to birds. One of ATOL’s objectives is to use trait terms related to industry-wide technical measurements to promote standardisation. This was realised through the adoption of PATO’s Entity-Quality formalism model^33^.

The Crop Ontology (CO)^79,80^ is a collection of crop-specific trait ontologies providing controlled vocabulary sets for 34 crop species, including a number of economically important plants such as wheat and sorghum. The effort aims to enhance interoperability in a field where many traits are crop-specific, have complex names and are normally captured in a free-text manner. The crop-specific trait ontologies were created following the Open Biomedical Ontologies (OBO) format standard to enable easy use and accessibility by biologists. The most common term relationships used in CO are “is_a”, “part_of”, “derived_from” and “has_a” and there are currently 7732 validated trait names.

The Fission Yeast Phenotype Ontology (FYPO)^81^ was constructed as a response to the need of Schizosaccharomyces pombe (S.pombe) researchers for phenotype annotation in a fission yeast database, a feature not present in PomBase or its predecessor GeneDB S.pombe. PomBase is a manually curated S. pombe comprehensive database. Similarly, GeneDB S.pombe is a manually constructed controlled vocabulary consisting of about 200 text descriptions. Terms in both of those S. pombe databases were not computable and did not allow for sharing or integration across species or with other databases. As a result, FYPO was created as a modular ontology drawing on terms from ontologies like PATO, GO and ChEBI to construct logical definitions using the Entity-Quality model in a pre-composed approach. The “entity” could be a whole or part of a cell, population of cells or an event, described by a “quality” usually captured by a PATO term. There are currently 20949 terms in FYPO organised along three main axes: abnormal and normal phenotypes, single cell and cell population phenotypes, or molecular function and biological process phenotypes.

Marine species traits^82^ are described in a number of databases and vocabularies such as WoRMS (The World Register of Marine Species), BIOTIC (The Biological Traits Information Catalogue), FishBase and SeaLifeBase. A broad but unstandardised terminology describes the biological attributes of marine life stages, reproduction, diet, body morphology as well as ecological traits. To promote standardisation, Costello et al.^82^ reviewed and prioritised ten top-level classifications of traits to form the foundation of a future marine species trait ontology. Semantic MediaWiki (SMW) was used to develop a hierarchy of traits, expressed in Simple Knowledge Organization System (SKOS)^83^, to enable a semantically interoperable ontology that can draw on other ontologies like PATO or the Environment Ontology (ENVO).

The World Spider Trait (WST)^84^ database is a centralised global open repository for curated data on spider traits. Any measurable phenotypic characteristic of an individual or taxon, like morphological, ecological, physiological or behavioural traits, are eligible for inclusion. It currently lists 12 categories and 223 traits, each with a description and abbreviation. Standardised terms are used to describe each trait in the WST to achieve semantic interoperability in order to fulfil a future goal of creating a hierarchical structure and detailed vocabularies for all traits and eventually an expertly developed ontology. Data in the WST is open-access, in a machine-readable format and was built to conform to the FAIR Guiding Principles for scientific data management.

## Conclusions

The Ontology of Biological Attributes (OBA) is a species-independent ontology with numerous links to other biological and biomedical ontologies that integrates widely used phenotype ontologies such as HPO. A scalable logical framework based on design patterns and templates allows the rapid curation of precisely defined terms which not only bridge the gap between low level characteristics (such as “weight” and “amount”) and reference ontologies such as the ChEBI (chemical entity) and Uberon (anatomy) ontologies, but also the currently wide chasm between quantitative “measurement” data such as GWAS and qualitative phenotyping data from clinical or model organism phenotyping activities. OBA is an active, evolving ontology that welcomes contributions and suggestions from the trait data community. In the near future, our goal is to integrate OBA more closely with clinical laboratory data (e.g. LOINC) and disease data (e.g. Mondo).

## Supplementary Materials

**S1:** All DOS-DP patterns used by OBA.

S1: All DOS-DP patterns used by OBA
| Pattern name | Description | Example |
| --- | --- | --- |
| attribute_location | This pattern describes an attribute of a specified entity that can be observed, measured or quantified in a specific location. | OBA:0000091 lysosomal lumen pH |
| chemical_role_attribute | This pattern describes an attribute of a chemical entity, where the chemical entity is a category of chemicals with a specific role. | OBA:0000079 hormone levels |
| chemical_role_attribute_location | This pattern describes an attribute of a chemical entity in a specified location, where the chemical entity is a category of chemicals with a specific role. | OBA:VT0003369 blood estrogen amount |
| disease age of onset | Age at which disease manifestations first appear. | OBA:2001000 age of onset of Alzheimer disease |
| entity_attribute | This pattern describes an attribute of a specific entity that can be observed, measured or quantified. | OBA:0002360 trochanter size |
| entity_attribute_location | This pattern describes an attribute of a specific entity that can be observed, measured or quantified in a specified location. | OBA:2020005 lysine in blood amount |
| entity_attribute_part_of | This pattern describes very general grouping traits. This should be used only in conjunction with morphology and quality attributes. | OBA:1000013 cytoskeleton morphology |
| process_attribute_location | This pattern describes an attribute of a biological process in some location. | OBA:2050069 serum lipase activity amount |
| response_to_chemical_stimulus_trait | This pattern describes a trait that affects the biological response to a chemical. | OBA:2040012 trait in response to morphine |
| response_to_chemical_with_role_stimulus | This pattern describes a trait that affects the biological response to a chemical, where the chemical is a category of chemicals with a specified role. | OBA:2040009 trait in response to endocrine disruptor |

**S2:**
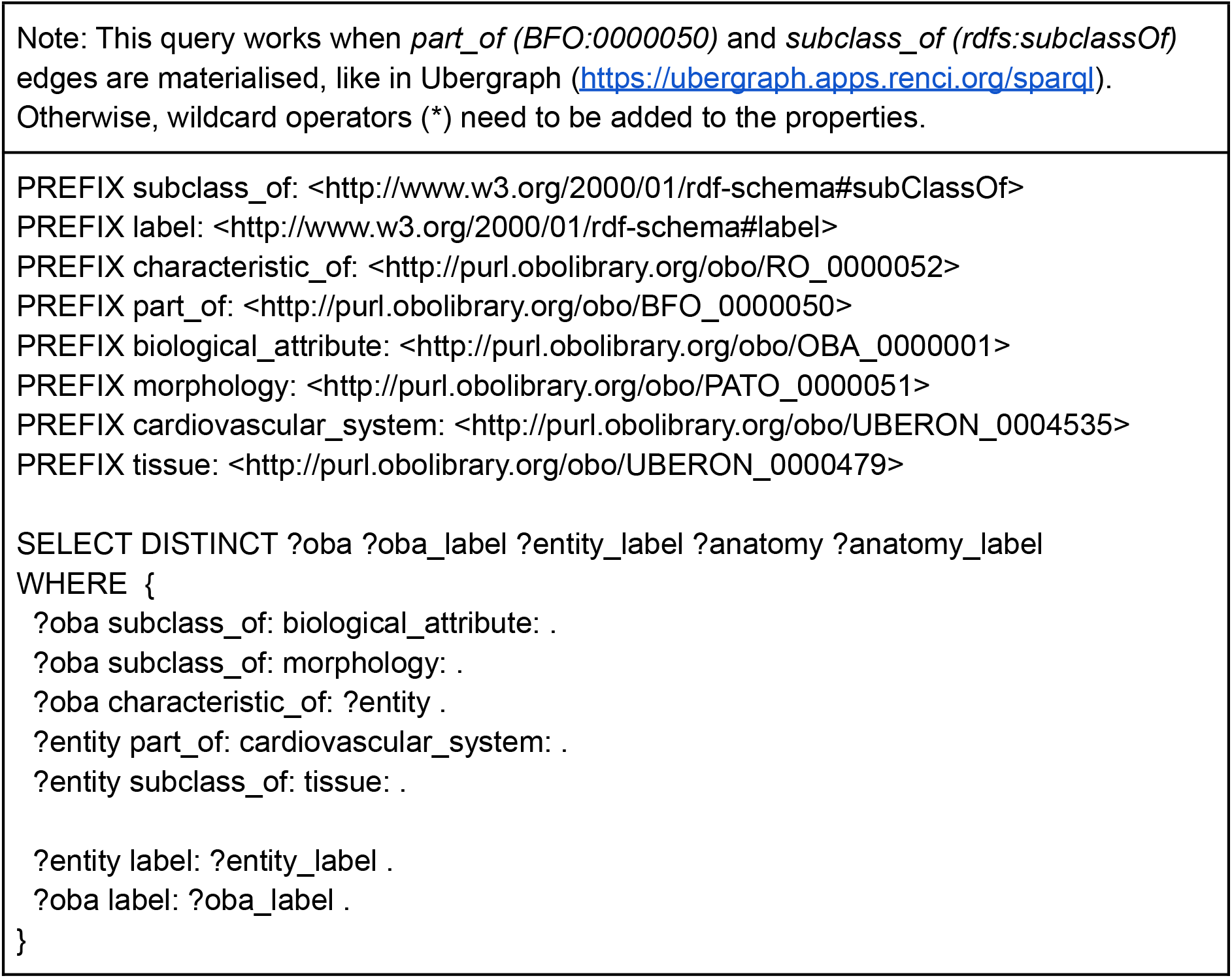
Supplementary query (SPARQL): Aggregate all data related to the morphology of the a part of the cardiovascular system.

